# Associations between canine temperament and salivary levels of cortisol and serotonin

**DOI:** 10.1101/2025.11.16.688729

**Authors:** Youngwook Jung, Yujin Song, Kayoung Yang, Kyongwon Yoo, Youngtae Heo, Minjung Yoon

## Abstract

Temperament influences canine behavior and helps determine a dog’s suitability as a companion or working animal. Although several temperament assessment tools exist, many rely on subjective evaluations, highlighting the need for scientifically validated methods. This study evaluated the validity of the Wesen temperament test by examining its association with physiological biomarkers—salivary cortisol and serotonin. Twenty-four dogs (11 females and 13 males) completed a modified Wesen test comprising seven subtests: Unconscious Confidence, Sociality, Noise Stability, Movement Stability, Desire for Play and Predation, Behavior in Stressful Situations, and Separation. Saliva samples were collected pre- and post-assessment, and cortisol and serotonin levels were measured using ELISA. Hormonal levels and temperament scores were analyzed using correlation, regression, and Kruskal-Wallis tests. Pre-assessment cortisol levels were negatively correlated with total average temperament score and several subtest scores. Post-assessment cortisol levels showed significant negative correlations with total average scores and all subtests. Changes in cortisol levels from pre- to post-assessment were negatively associated with temperament scores. Dogs with higher temperament scores exhibited significantly higher serotonin levels than those with lower scores. These findings support the utility of temperament assessments validated through physiological markers and provide novel evidence linking canine temperament with endocrine function.

## 1. Introduction

The temperament of dogs is a key factor in determining their suitability as companion animals [1]. Additionally, temperament plays a critical role in working contexts, such as guiding [2], detection [3], and rescue operations [4]. Therefore, selecting dogs with appropriate temperaments for specific purposes is important. Temperament assessment can help predict a dog’s future performance and suitability [5]. Several tools have been developed for this purpose, including the Canine Behaviour Assessment and Research Questionnaire (C-BARQ) [6], the In-For Training test [7], and the Wesen test [8]. These assessment tools differ in their evaluation methods: Some rely on owners’ reports, while others require trained professionals. This variability raises concerns about potential subjectivity [7]. Thus, several studies have examined the validity of temperament assessments by comparing their scores with physiological biomarkers such as heart rate, respiratory rate, and hormone levels [9–11]. Among these biomarkers, hormones—particularly cortisol and serotonin—have emerged as key indicators owing to their close association with temperament and behavior.

Cortisol is secreted in response to stress or negative emotional states [12]. It is commonly used as a biomarker of stress and overall well-being in dogs. Recent research has focused on the relationship between cortisol and temperament. Rosado and colleagues reported higher cortisol levels in aggressive dogs than in non-aggressive individuals [13]. Basal cortisol levels have also been linked to fear-related behaviors, such as panting and tongue protrusion [14]. Additionally, cortisol levels vary with the degree of sociality [15] and have been associated with touch sensitivity [16] and anxiety towards strangers [17]. Collectively, these findings indicate that cortisol is involved in several aspects of canine temperament, including aggression, fear responses, and social interaction. Therefore, cortisol serves as an indicator of temperament-related traits beyond its traditional role in stress response.

Cortisol has been measured in several biological matrices, including plasma, urine, feces, hair, feathers, and saliva [14, 18–20]. Blood cortisol is considered a reliable indicator of acute stress responses. However, because blood sampling is invasive, it has become less frequently used in recent studies. In contrast, saliva sampling has gained popularity as a noninvasive method that reduces stress in animals. Steroid hormones can diffuse rapidly into saliva from the bloodstream owing to their molecular properties [21]. Moreover, salivary cortisol reliably reflects plasma cortisol levels in dogs [22]. Therefore, we selected saliva sampling as the method for cortisol measurement.

Serotonin plays a well-established role in temperament and behavior, including aggression, anxiety, and social interactions. Numerous studies have reported substantial differences in serotonin levels between aggressive and non-aggressive dogs [13, 23–25]. Vermeire and colleagues further reported that serotonin receptors are expressed in cortical regions of the brain and are associated with anxiety disorders in dogs [26]. Associations between serotonin levels and dogs’ social behaviors toward humans have also been observed [27]. Collectively, these findings suggest that serotonin is a biomarker for temperament assessment in dogs.

In dogs, serotonin has predominantly been measured in blood samples [25, 28–30]. However, the invasiveness of blood sampling may limit its use in temperament monitoring. Therefore, noninvasive alternatives are needed. Saliva sampling has become increasingly popular because it minimizes stress in animals. In humans, although salivary serotonin is not directly correlated with serotonin levels in cerebrospinal fluid or platelets, it provides a reliable measure of peripheral serotonin levels [31, 32]. Furthermore, salivary serotonin levels have been associated with indicators of happiness, mood, and depressive disorders [33–35]. On this basis, saliva sampling was adopted as the method for measuring serotonin levels in this study.

This study aimed to validate a temperament assessment tool by measuring salivary cortisol and serotonin levels. Specifically, we (i) examined correlations between temperament scores and cortisol and serotonin levels and (ii) compared hormone levels between high- and low-scoring groups, as determined by the temperament assessment. We hypothesized that salivary hormone levels would correlate with temperament traits assessed by the Wesen test, thereby providing physiological evidence for the convergent validity of the test. The findings of this study may provide a scientific basis for the use of temperament assessments in dogs.

## 2. Materials and methods

### 2.1. Subjects

Twenty-four dogs (eleven females and thirteen males) participated in the study, including seven mixedbreeds, six Beagles, four Border Collies, two Labrador Retrievers, two Malinois, one Pomeranian, one French Bulldog, and one Shepherd. The mean age was 5.13 ± 3.06 years. The Beagles were bred for experimental use, whereas the remaining dogs were owned by private individuals or professional trainers. All experimental procedures were conducted in accordance with institutional ethical guidelines and were approved by the Animal Experimentation Ethics Committee of Kyungpook National University (approval number: 2024-0531).

### 2.2. Temperament assessment

To minimize stress associated with transportation, temperament assessments were conducted at three locations near the dogs’ original environments. The assessment was led by an experienced international dog show judge using a slightly modified version of the Wesen test. An overview of the temperament test process is presented in Fig 1. The Wesen test (Wesensüberprüfung) was originally developed for German Shepherds aged 9–13 months [36], but its effectiveness and adaptability have led to its broad application across breeds and age groups [37]. The modified version used in this study consisted of seven subtests [5, 8, 36]:

1. Unconscious Confidence: Attention, fear, confidence, interest, and relaxation were assessed during a physical examination that included inspection of the teeth, testes (for males), and a full-body check by the judge.
2. Sociality: Responsiveness to the handler’s cues, confidence, cheerfulness, affinity, and energy were assessed during encounters with a group of strangers and an unfamiliar dog.
3. Noise Stability: Calmness, relaxation, fear, and confidence were assessed in response to auditory stimuli produced by a motor, metal chain, and whip.
4. Movement Stability: Dogs were guided by handlers to sit and walk on an unstable pallet and a series of connected tables. Relaxation, stability, fear, and activity were evaluated.
5. Desire for Play and Predation: Dogs played with a toy or treat provided by both the handler and a stranger and were tasked with finding an object hidden under a box. Activity, persistence, affinity, and interest were evaluated.
6. Behavior in Stressful Situations: Dogs played with a toy provided by the handler. Thereafter, the toy was hidden to prompt a search. During the search, a sudden loud noise was produced by dropping a metal chain to simulate a stress-inducing stimulus. Activity, confidence, persistence, interest, and reactions to the noise were evaluated.
7. Separation: Dogs were tied to a fixed object and left alone for 5 min, followed by an encounter with a stranger. Anxiety, relaxation, and affinity were evaluated.

**Fig 1.**
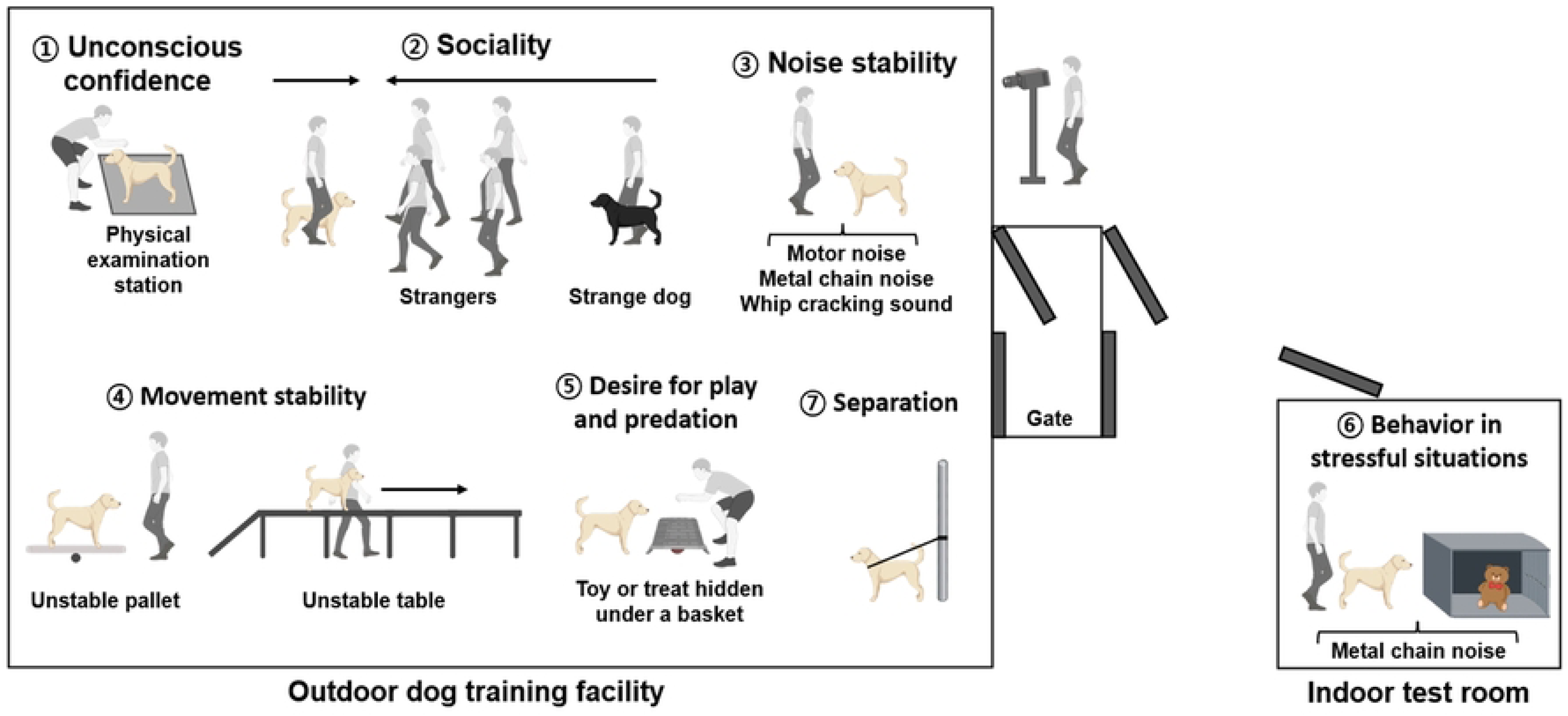
Schematic representation of the temperament assessment test process (created using BioRender.com)

Each subtest comprised two to four categories, scored on a five-point scale (1 = lowest; 5 = highest). The judge evaluated all reactions according to standardized guidelines. During the Unconscious Confidence subtest, the judge held the leash; in all remaining subtests, the dog was handled either by the handler or a stranger. The entire temperament assessment was video-recorded by a designated observer. All subtests were conducted at an outdoor dog-training facility, except for the Behavior in Stressful Situations subtest, which took place in an indoor test room.

### 2.3. Sample collection

To analyze hormonal levels, saliva samples were collected from each dog before and after temperament assessment by an experimenter not involved in judging. Samples were collected using a swab device and storage tube (Salimetrics, Carlsbad, CA, USA) by gently placing the swab under the tongue and in the cheek pouch. During transport to the laboratory, the samples were kept in an icebox at 4 ℃ to maintain integrity. Upon arrival, saliva was centrifuged at 1,000 × *g* for 2 min at 4 ℃ and stored at −80 ℃ until analysis.

### 2.4. Hormone analysis

Hormone concentrations were measured using commercial enzyme-linked immunosorbent assay (ELISA) kits. Samples were analyzed in duplicate without prior extraction or dilution. Optical density was measured using a microplate reader (Tecan, Männedorf, Switzerland), and concentrations were calculated based on a four-parameter logistic standard curve.

#### 2.4.1. Cortisol ELISA

Salivary cortisol levels before and after temperament assessment were measured using a Cortisol ELISA kit (1-3002; Salimetrics) with a sensitivity of 0.018 μg/dL. The intra- and inter-assay coefficients of variation (CVs) were 1.55% and 3.57%, respectively. Optical density was measured at 450 nm following the manufacturer’s instructions.

#### 2.4.2. Serotonin ELISA

Owing to insufficient saliva volume, sixteen samples that met the minimum quantity required for ELISA were included in the serotonin analysis. Salivary serotonin levels before temperament assessment were determined using a Serotonin ELISA kit (ADI-900-175; Enzo Life Sciences, Farmingdale, NY, USA) with a sensitivity of 0.293 ng/mL. The intra- and inter-assay CVs were 7.08% and 14.97%, respectively. Optical density was measured at 405 nm following the manufacturer’s protocol.

### 2.5. Statistical analysis

Statistical analyses were performed using IBM SPSS Statistics (version 27; IBM, Armonk, NY, USA), and data visualization was conducted using GraphPad Prism (version 9; GraphPad Software, San Diego, CA, USA). Data normality was assessed using the Shapiro–Wilk test. Sex differences in hormone concentrations were evaluated using the Mann–Whitney *U* test. Variations in hormone concentrations among dogs bred for experimental use, privately owned dogs, and dogs raised by professional trainers were evaluated using the Kruskal–Wallis test. Spearman’s rank correlation coefficient (ρ) was used to examine relationships between hormone concentrations and temperament scores. Additionally, both Spearman’s ρ and linear regression analyses were used to evaluate associations between changes in cortisol concentrations and total temperament scores. The Kruskal–Wallis test was also used to compare hormone concentrations across groups categorized by temperament scores. Hormone concentrations and temperament scores are presented as mean ± standard error of the mean. Statistical significance was set at *p* < 0.05, and values between 0.05 and 0.1 were considered indicative of a trend.

## 3. Results

### 3.1. Correlation between cortisol levels and temperament assessment scores

To examine potential sex effects, hormone concentrations were compared between females and males. No significant differences in cortisol or serotonin levels were observed between sexes or among dog populations. Therefore, data were pooled across sex and population for subsequent analyses. Correlations between salivary cortisol levels and temperament assessment scores are presented in Table 1. Pre-assessment cortisol levels were negatively correlated with total average scores (*p* = 0.036). Negative correlations were also observed between pre-assessment cortisol levels and several subtests, including Unconscious Confidence (*p* = 0.006), Sociality (*p* = 0.035), Noise Stability (*p* = 0.003), and Separation (*p* = 0.006). No significant correlations were observed between pre-assessment cortisol levels and other subtests, including Movement Stability (*p* = 0.291), Desire for Play and Predation (*p* = 0.801), and Behavior in Stressful Situations (*p* = 0.126). Post-assessment cortisol levels were negatively correlated with all subtest scores as well as with the total average scores (*p* < 0.001).

### 3.2. Correlation between changes in cortisol levels and temperament assessment scores

The correlation between changes in cortisol levels (from pre- to post-assessment) and total average scores is illustrated in Fig 2. Cortisol levels were negatively correlated with total average scores (*p* = 0.008). The coefficient of determination (*r*²) was 0.259 (*p* = 0.011), and the adjusted *r*² was 0.225.

**Fig 2.**
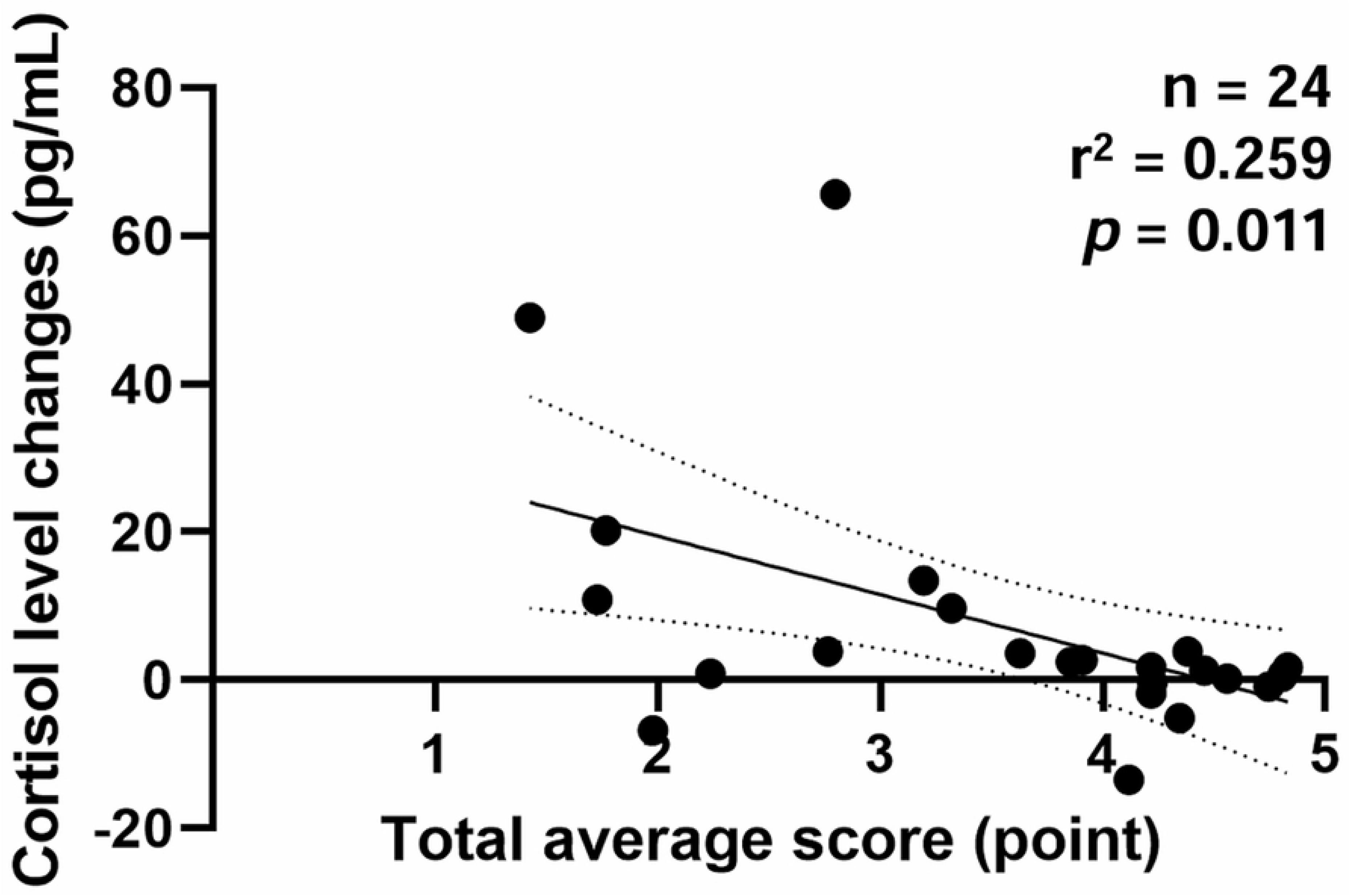
Linear regression between changes in cortisol levels (pre- to post-assessment) and total average temperament assessment scores. The analysis indicates a negative correlation (Spearman’s ρ = −0.526, *p* = 0.008) with an *r*² of 0.259 (*p* = 0.011) and an adjusted *r*² of 0.225.

### 3.3. Comparison of cortisol levels among three groups

Cortisol levels tended to increase from pre- to post-assessment (8.339 ± 1.352 pg/mL vs. 15.184 ± 3.682 pg/mL; *p* = 0.062). To further investigate these differences, dogs were categorized into three scoring groups—high (n = 8), medium (n = 8), and low (n = 8)—based on their total average temperament assessment scores (Table 2). These groups differed significantly in their total average scores (*p* < 0.01). Pre-assessment cortisol levels tended to be lower in the high-scoring group compared with the low-scoring group (*p* = 0.058; Fig 3). Post-assessment cortisol levels were higher in the low-scoring group than in both the high-scoring (*p* = 0.005) and medium-scoring (*p* = 0.012) groups. Within the low-scoring group, post-assessment cortisol levels also tended to be higher than pre-assessment levels (*p* = 0.059).

**Fig 3.**
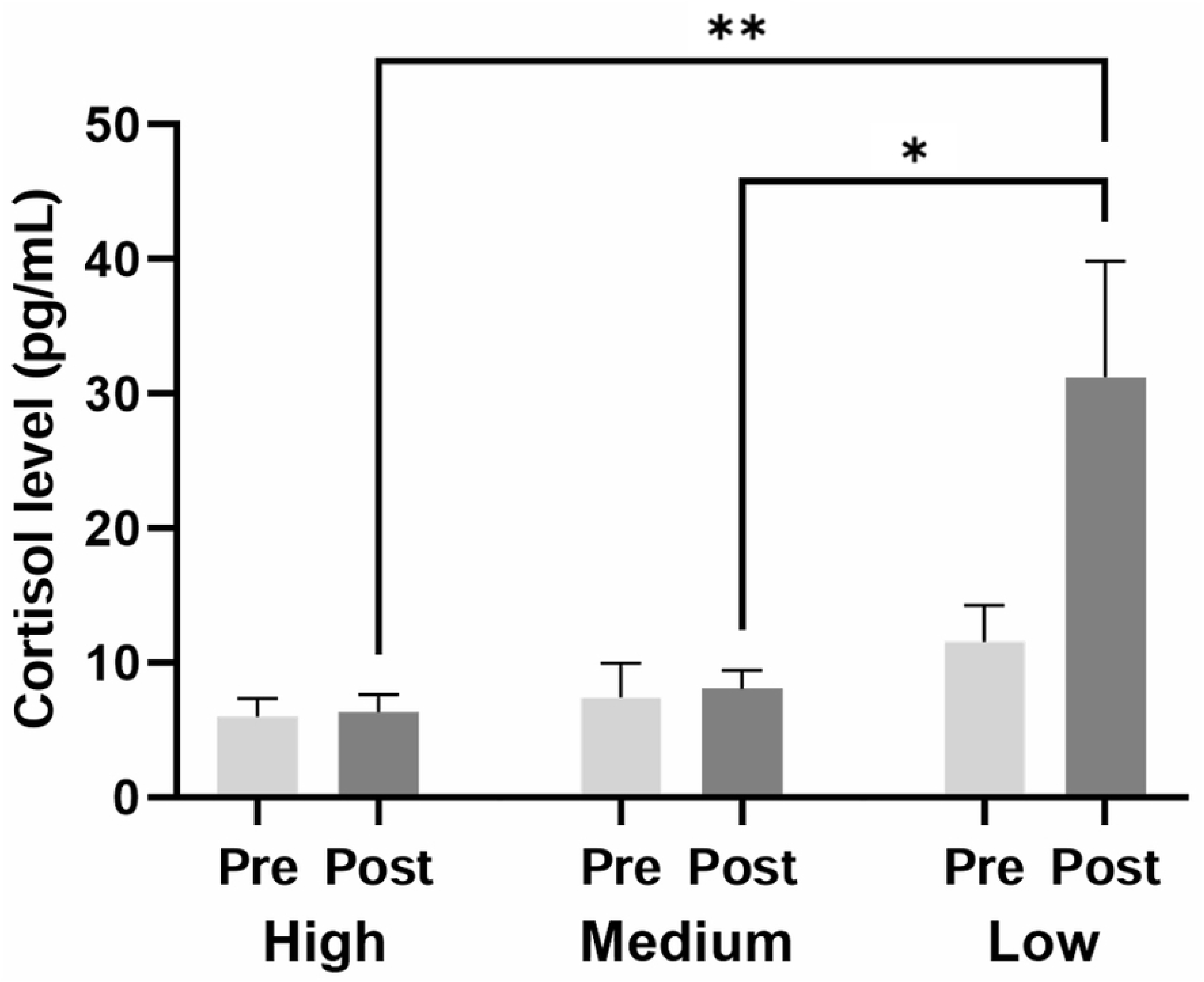
Salivary cortisol levels in high-, medium-, and low-scoring groups. Cortisol levels were 5.996 ± 1.258 pg/mL (pre) and 6.295 ± 1.283 pg/mL (post) in the high-scoring group, 7.441 ± 2.379 pg/mL (pre) and 8.044 ± 1.296 pg/mL (post) in the medium-scoring group, and 11.581 ± 2.535 pg/mL (pre) and 31.213 ± 8.075 pg/mL (post) in the low-scoring group. Pre and Post indicate samples collected before and after temperament assessment, respectively. ** *p* < 0.01, * *p* < 0.05

### 3.4. Correlation between serotonin levels and temperament assessment scores

Correlations between salivary serotonin levels and temperament assessment scores are summarized in Table 1. A trend toward a positive correlation was observed between serotonin levels and the total average scores, although this result was not statistically significant (*p* = 0.064). In contrast, a significant positive correlation was identified between serotonin levels and the Movement Stability subtest (*p* = 0.009). No significant correlations were observed for the remaining subtests, including Unconscious Confidence (*p* = 0.129), Sociality (*p* = 0.118), Noise Stability (*p* = 0.165), Desire for Play and Predation (*p* = 0.108), Behavior in Stressful Situations (*p* = 0.277), and Separation (*p* = 0.159).

**Table 1.** Correlations between temperament assessment scores and salivary cortisol and serotonin levels.

|  |  | Total<br>average<br>scores | Unconscious<br>Confidence | Sociality | Noise<br>Stability | Movement<br>Stability | Desire for<br>Play and<br>Predation | Behavior in<br>Stressful<br>Situations | Separation |
| --- | --- | --- | --- | --- | --- | --- | --- | --- | --- |
| Cortisol<br>(Pre) | Spearman's $\rho$ | -0.430* | -0.544** | -0.432* | -0.586** | -0.225 | -0.054 | -0.321 | -0.540** |
| | $p$-value | 0.036 | 0.006 | 0.035 | 0.003 | 0.291 | 0.801 | 0.126 | 0.006 |
| Cortisol<br>(Post) | Spearman's $\rho$ | -0.714** | -0.540** | -0.557** | -0.705** | -0.617** | -0.497* | -0.598** | -0.817** |
| | $p$-value | <0.001 | 0.006 | 0.005 | <0.001 | 0.001 | 0.013 | 0.002 | <0.001 |
| Serotonin | Spearman's $\rho$ | 0.474<sup>#</sup> | 0.396 | 0.406 | 0.364 | 0.633** | 0.417 | 0.289 | 0.369 |
| | $p$-value | 0.064 | 0.129 | 0.118 | 0.165 | 0.009 | 0.108 | 0.277 | 0.159 |

\*\* $p < 0.01$ , \* $p < 0.05$ , <sup>#</sup> $0.05 < p < 0.1$
Pre and Post indicate samples collected before and after temperament assessment, respectively.

### 3.5. Comparison of serotonin levels among three groups

To compare serotonin levels among groups, dogs were categorized into high- (n = 6), medium-(n = 5), and low-scoring (n = 5) groups based on their total average temperament assessment scores (Table 2). Significant differences in the total average scores were observed among the groups (*p* < 0.01). Serotonin levels in the high-scoring group were significantly higher than those in the low-scoring group (*p* = 0.028; Fig 4).

**Table 2.** Average temperament assessment scores of each group for cortisol and serotonin analyses.

|  | High | Medium | Low |
| --- | --- | --- | --- |
| Cortisol | 4.618 ± 0.068<sup>a</sup> | 3.937 ± 0.109<sup>b</sup> | 2.237 ± 0.206<sup>c</sup> |
| Serotonin | 4.368 ± 0.093<sup>a</sup> | 3.360 ± 0.163<sup>b</sup> | 1.828 ± 0.121<sup>c</sup> |

Different superscripts indicate a significant difference ( $p < 0.01$ ).

**Fig 4.**
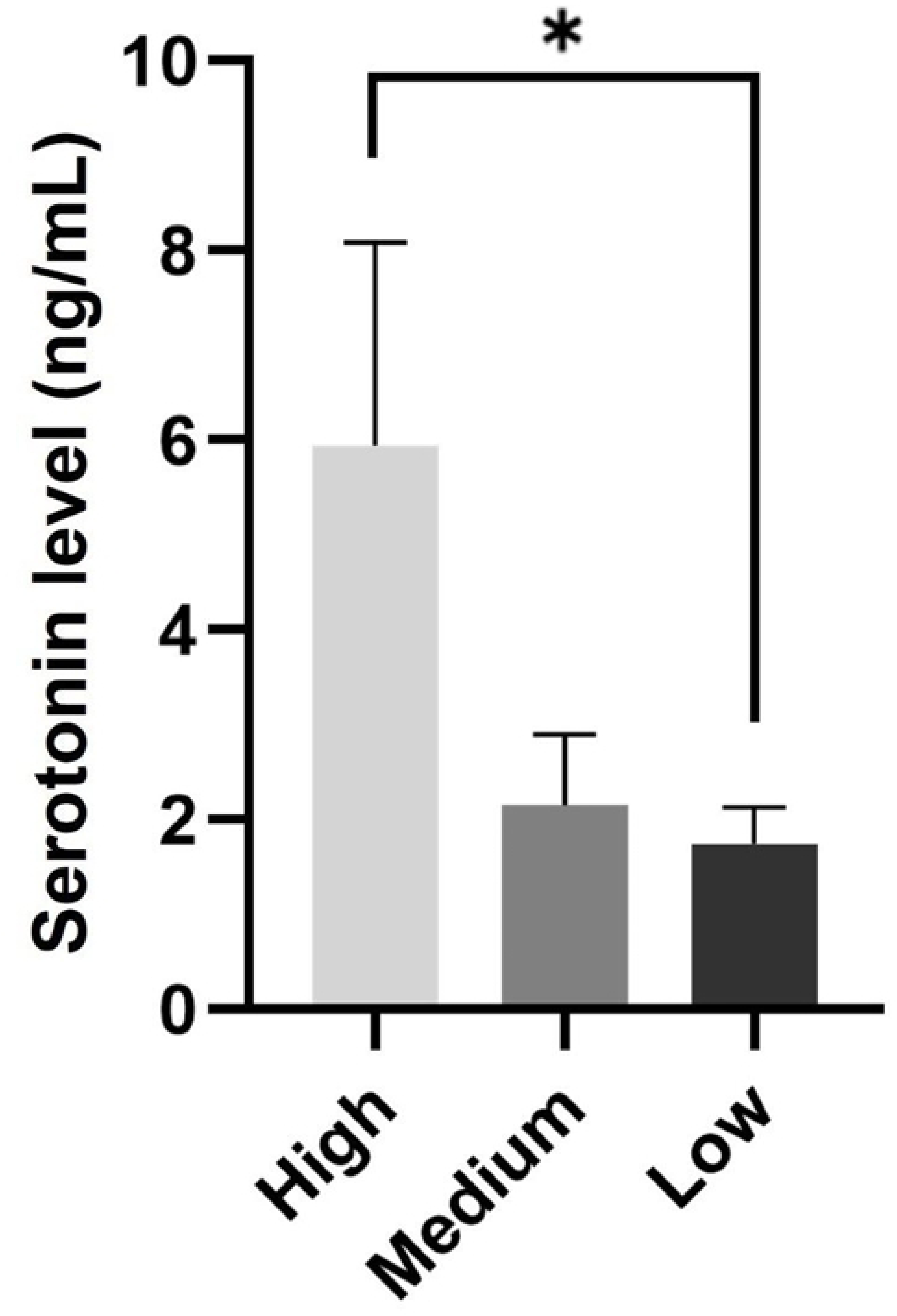
Salivary serotonin levels in high-, medium-, and low-scoring groups. Serotonin levels were 5.943 ± 1.958 ng/mL in the high-scoring group, 2.154 ± 0.666 ng/mL in the medium-scoring group, and 1.742 ± 0.348 ng/mL in the low-scoring group. * *p* < 0.05

## 4. Discussion

In this study, we validated a temperament assessment based on the Wesen test by incorporating hormonal markers as physiological indicators. Establishing such physiological evidence strengthens the foundation for accurately evaluating canine temperament. A reliable assessment tool can help determine a dog’s suitability as a companion animal or for specific working purposes. Moreover, physiologically validated assessments may contribute to policies and legislation that promote the safe selection and responsible adoption of dogs with appropriate temperaments.

We examined the associations between temperament assessment scores and salivary cortisol levels. Pre-assessment cortisol levels were negatively correlated with the total average score and several subtests. Post-assessment cortisol levels showed negative correlations with the total average score and all subtests. These findings provide physiological evidence supporting the convergent validity of the Wesen test. Lower cortisol levels, an established biomarker of acute stress and arousal in dogs, were associated with greater emotional stability. Our results align with previous studies reporting associations between cortisol levels and behavioral traits. Rayment and colleagues reported a slight positive correlation between cortisol levels and C-BARQ traits such as separation-related problems and touch sensitivity [16]. Similarly, McPeake and colleagues observed a negative correlation between post-assessment salivary cortisol levels and frustration-coping ability [10]. In a study investigating the relationship between long-term stress and temperament, Roth and colleagues discovered that hair cortisol levels were positively correlated with stranger-directed aggression and chasing traits, both assessed using the C-BARQ [9]. Rossi and colleagues also reported that lower cortisol levels were associated with more frequent play-soliciting and longer durations of exploratory behavior [38]. Collectively, these findings suggest that cortisol levels reflect behavioral traits related to stress responsiveness and emotional regulation across several temperament assessment tools.

In this study, we identified a relationship between temperament scores and changes in salivary cortisol levels. Dogs with higher temperament scores exhibited more stable physiological responses. The negative correlation between cortisol changes and Wesen test scores suggests that behaviorally well-adjusted dogs may be more resilient to stress. These findings support the convergent validity of the Wesen test in assessing emotional stability and stress reactivity in dogs. Similar findings have been reported in other studies. McPeake and colleagues reported significant correlations between scores on the Canine Frustration Questionnaire and changes in salivary cortisol levels [10]. Similarly, the startling test has been validated through its association with physiological indicators such as cortisol levels and heart rate [11]. Another study found that high sociability, as measured by behavioral assessment tests, was associated with smaller variations in cortisol levels [39]. Overall, these findings underscore the consistent relationship between canine temperament traits—across several temperament assessment tools—and physiological responses related to stress.

Additionally, we compared cortisol levels among three groups classified by temperament assessment scores. Pre-assessment cortisol levels tended to be lower in the high-scoring group than in the low-scoring group, consistent with reports that shelter dogs with more sociable temperaments exhibit lower baseline cortisol levels [40]. Likewise, dogs engaging in more positive interactions generally exhibit lower cortisol levels [41]. Post-assessment cortisol levels were significantly higher in the low-scoring group than in the medium- and high-scoring groups, indicating greater stress reactivity among dogs with lower temperament scores. Consistent with this finding, Shin and Shin reported that more sociable dogs exhibited lower cortisol levels following behavioral assessment than less sociable dogs [39]. These findings suggest that dogs with more stable or socially adaptive temperaments are less reactive to testing situations and experience lower stress levels. In the low-scoring group, cortisol levels tended to increase after the assessment, reflecting the overall trend observed in this study. Collectively, these results indicate that both basal and reactive cortisol levels reflect underlying temperament traits, with socially adaptive dogs exhibiting lower physiological stress responses in routine and test-related situations.

Interestingly, Lensen and colleagues reported that salivary cortisol and chromogranin A (CgA) levels measured 10 min after a behavioral test were positively associated with high trainability and negatively associated with stranger-directed fear, respectively, in adult dogs [42]. However, these associations varied depending on the timing of sample collection. Cortisol levels measured 40 min after the test were associated with high energy, whereas CgA levels were related to low excitability, both of which may be considered undesirable traits. These findings highlight the importance of sampling time when interpreting stress-related biomarkers.

We also analyzed the relationship between temperament assessment scores and salivary serotonin levels. Although overall temperament scores did not show a statistically significant correlation with serotonin levels, a positive trend was observed. This finding suggests that behavioral indicators of emotional stability are partly associated with serotonergic activity. The Movement Stability subtest, which evaluates traits such as relaxation, stability, and activeness, showed a significant positive correlation with serotonin levels. This finding supports the convergent validity of the subtest, indicating that higher serotonin levels are associated with calmer and more regulated behaviors. These results are consistent with previous findings on serotonin–behavior relationships in dogs. Rayment and colleagues reported a negative trend between serotonin levels and C-BARQ traits related to separation-related problems and touch sensitivity [16]. Similarly, another study identified a weak linear correlation between social behavioral scores and serotonin levels in shelter dogs [27]. Wright and colleagues also found that higher impulsivity scores on the Dog Impulsivity Assessment Scale (DIAS) were associated with lower serotonin levels [43]. Taken together, these studies and our findings suggest that serotonin levels are associated with several positive behavioral characteristics across diverse behavioral assessment tools. This highlights the potential of serotonin as a physiological marker of emotional and behavioral regulation in dogs.

We also compared serotonin levels among three groups classified by temperament scores. Dogs in the high-scoring group exhibited significantly higher serotonin levels than those in the low-scoring group. This finding aligns with numerous studies reporting that higher serotonin levels are associated with lower aggression and anxiety in dogs [27, 30, 44–46]. Collectively, these findings provide physiological evidence supporting the reliability of the temperament assessment used in this study. However, one study reported no significant differences in serotonin levels between groups classified by factors assessed through the C-BARQ or DIAS [16]. This inconsistency may stem from methodological differences. Unlike the temperament assessment in our study, which involved direct behavioral observation, both the C-BARQ and DIAS rely on owner-reported data. Although these tools are validated, their outcomes can be influenced by individual variation in owner perception and experience. Moreover, previous studies suggest that factors such as learning history, environmental conditions, and breed characteristics also influence test outcomes [16]. Therefore, these factors should be considered when interpreting associations between behavioral assessments and physiological markers.

Although this study yielded several noteworthy findings, certain limitations should be considered when interpreting the results. Owing to constraints in salivary sampling, fewer samples were available for serotonin analysis, resulting in a smaller dataset compared with that of cortisol. To improve reliability and generalizability, future studies should include a larger number of dogs. Previous research has shown that salivary stimulants do not significantly affect hormonal measurements [47]. Thus, their use may be considered in future studies to increase saliva yield. In this study, no significant differences in hormonal levels were observed between groups classified by sex or living environment. Nevertheless, these factors may influence hormonal responses under different conditions or study designs and should be considered in future investigations. Cobb and colleagues reported that variables such as breed and body weight have minimal effects on salivary cortisol levels [47]; however, they also found that intact females exhibited significantly higher cortisol levels than intact males and neutered dogs. Additionally, living environment influenced cortisol levels, and puppies under 6 months of age exhibited significantly lower cortisol levels than older dogs. Similarly, Sandri and colleagues found that cortisol levels were associated with body size and housing conditions [48]. Collectively, these findings suggest that biological and environmental factors, including body size, sex, reproductive status, age, and housing conditions, influence basal cortisol levels. Future research should explore how these factors interact with temperament assessment outcomes, such as those derived from the Wesen test, to clarify their combined effects on physiological stress responses in dogs. Although cortisol is a well-established biomarker of stress, future studies should incorporate other physiological indicators to improve the robustness of findings [49]. Heart rate is one such parameter. For example, Robinson and colleagues reported that heart rate correlated with several behavioral test batteries [50]. Moreover, King and colleagues found a strong correlation between heart rate and startling test scores [11]. Incorporating validated physiological markers and examining their interrelationships may enhance the reliability and robustness of future findings.

In conclusion, this study validated a canine temperament assessment tool by examining its associations with cortisol and serotonin. Significant correlations were observed between temperament assessment scores and cortisol levels, and a potential association was observed with serotonin levels. Furthermore, both hormonal markers differed among groups categorized by temperament scores. These results support the utility of temperament assessments validated through physiological measures and contribute to a better understanding of the relationships among canine temperament, behavior, and endocrine function.

## Acknowledgements

The authors would like to thank Dr. Yeonju Choi (Malgill Healing and Leadership, Republic of Korea) and Yubin Song, Taelang Kim, Jaewoo Choi, and Junseo Ko (Kyungpook National University, Republic of Korea) for their assistance with the temperament assessment. We also extend our gratitude to Dr. Heejun Jung (Korea Polytechnics, Republic of Korea) and Junyoung Kim, Gaeun Jeong, and Suhyun Kim (Kyungpook National University, Republic of Korea) for their support.

## Author contributions

**Conceptualization**: Youngwook Jung, Kyongwon Yoo, Youngtae Heo, Minjung Yoon

**Data curation**: Youngwook Jung

**Formal analysis**: Youngwook Jung

**Funding acquisition**: Kayoung Yang, Kyongwon Yoo, Youngtae Heo, Minjung Yoon

**Investigation**: Youngwook Jung, Yujin Song, Kayoung Yang, Kyongwon Yoo, Youngtae Heo, Minjung Yoon

**Methodology**: Youngwook Jung, Yujin Song, Kayoung Yang, Kyongwon Yoo, Youngtae Heo, Minjung Yoon

**Project administration**: Minjung Yoon

**Resources**: Kayoung Yang, Kyongwon Yoo, Youngtae Heo, Minjung Yoon

**Software**: Youngwook Jung

**Supervision**: Kayoung Yang, Kyongwon Yoo, Youngtae Heo, Minjung Yoon

**Validation**: Youngwook Jung

**Visualization**: Youngwook Jung

**Writing – original draft**: Youngwook Jung

**Writing - review & editing**: Youngwook Jung, Minjung Yoon

